# AncientMetagenomeDir dating metadataset highlights need for standardised radiocarbon reporting in ancient DNA

**DOI:** 10.64898/2026.02.06.704039

**Authors:** Katherine Hearne, Diāna Spurīte, Alexander Hübner, Biancamaria Bonucci, Yuejiao Huang, Bjørn Peare Bartholdy, Piotr Rozwalak, Lorena Becerra-Valdivia, Maxime Borry, Davide Bozzi, Anastasia Brativnyk, Jodie Brunt, Nihan D. Dagtas, Toni de-Dios, Cameron Ferguson, Jasmin Frangenberg, Benjamin Guinet, Magdalena Haller-Caskie, Reed Harder, Iseult Jackson, Chenyu Jin, Marcel Keller, Arthur Kocher, Ian Light-Maka, Maria Lopopolo, Frederik Lutz, Megan Michel, Denis Jorge Nunes Yamunaque, Kadir Özdogan, Zoé Pochon, Darío A. Ramirez, Maximilian L. Schumacher, Gabriel Yaxal Ponce-Soto, Nikolaos Psonis, Tom Richtermeier, Lennart Schreiber, Pooja Swali, Cathy Ngọc Hân Trần, Irina M. Velsko, Christina Warinner, Anna E. White, Giulia Zampirolo, James A. Fellows Yates, Aida Andrades Valtueña

## Abstract

Ancient DNA is a valuable data source for the understanding of our past. However, to effectively interpret this data, it is essential to know the age of the samples from which the DNA is obtained. Although the field of palaeogenomics has been recognised for its robust open data sharing practices, dating information associated with analysed samples is not reported consistently across palaeogenomic studies, nor is it included as metadata in most genetic data repositories. Here, we describe the addition of standardised precise dating information for ancient microbial genomes into the AncientMetagenomeDir metadata repository of published ancient metagenomic samples. This extension currently includes dating information for over 700 ancient microbial genomic datasets, of which 333 are dated using historical, contextual, or stratigraphic methods, and 405 are radiocarbon dated. We quantitatively assess the quality of radiocarbon date reporting and find that, despite established reporting conventions, radiocarbon dating information is often reported inconsistently across ancient metagenomic studies. This new resource provides ancient microbial researchers with standardised dating information that facilitates more accurate and consistent analysis of metagenomic sequencing data. The dataset also highlights the need for greater standardisation of radiocarbon date reporting in original publications in order to allow effective reuse of this and future ancient microbial data.

## Background & Summary

Understanding the temporal context of a sample is essential to contextualise the findings from the palaeo– and archaeosciences. One of the most important methods for generating absolute dates from organic materials is radiocarbon dating. A radiocarbon age is derived from measurements on the radioactive decay of the carbon isotope ^14^C^1–3^, representing a proxy for the passage of time rather than a true calendar date. Radiocarbon dates are therefore calibrated to estimate calendar ages using standard atmospheric and marine curves, which account for variations in atmospheric radiocarbon production and are routinely updated^4–6^. Non-atmospheric reservoir effects^7,8^ and pretreatment methodologies for sample decontamination^9^ have an important impact on date accuracy. For example, the consumption of marine foods can result in erroneously older ages if not properly corrected^10^, and differences in pretreatment protocols can yield estimates that diverge by millennia for highly contaminated or ‘old’ (e.g., >25,000 yr BP bone collagen samples^11–17^). Adequate reporting of chronometric data, including uncalibrated ages (‘raw’ radiocarbon measurements – 14C Age [yr BP]), pretreatment methodology, stable isotope data, and radiocarbon quality control parameters (e.g., collagen yield, %C, C:N values) are key for temporal reliability. Reporting guidelines have been developed within the radiocarbon community^4,5^ to ensure the robustness, reproducibility, and reuse of radiocarbon data. However, compliance with these reporting standards within palaeo– and archaeo-science publications is often suboptimal^6–8^.

Sample age information is critical to contextualise palaeogenomic findings regarding past populations, such as population movement and kinship pedigrees^9–12^, modelling the presence of different strains or lineages of pathogens through time^13–17^, or understanding ancient ecological dynamics^18–20^. However, radiocarbon dating information is tedious to extract from article PDFs in the published literature, and in the absence of a centralised database for radiocarbon dates or a ‘gold-standard’ file format (e.g., equivalent to a FASTQ^21^ or BAM^22^ file in genetics), it is not possible to systematise the process. Researchers do have the ability to annotate genetic data with dating information during data upload to online repositories, but as this relies on the use of free text note fields rather than dedicated fields or checklists, it is often not done. In an analysis of 42 aDNA studies with human, animal, and plant genomic data uploaded to public databases, Bergström (2024) found that 83% of the data lacked sample age information in their uploaded metadata. While ancient genomics has previously been celebrated for a strong culture of openly sharing sequencing data^23,24^, associated metadata does not reach the same quality.

Initiatives such as AncientMetagenomeDir^25,26^ have been established to improve compliance of ancient metagenomic sequencing metadata, in particular ancient microbial genomes and microbiomes, with FAIR (Findable, Accessible, Interoperable, and Reusable) principles^27^. AncientMetagenomeDir provides structured and standardised tab-separated value (TSV) tables of annotated ancient metagenomic samples and library information from published studies, facilitating efficient searching and retrieval of published data. Since its initial release in 2021 (v20.09), AncientMetagenomeDir has undergone several major updates, increasing the number of incorporated publications from 87 to 216, and expanding the total number of samples from 1,024 to 3,222 (Figure 1).

**Figure 1:**
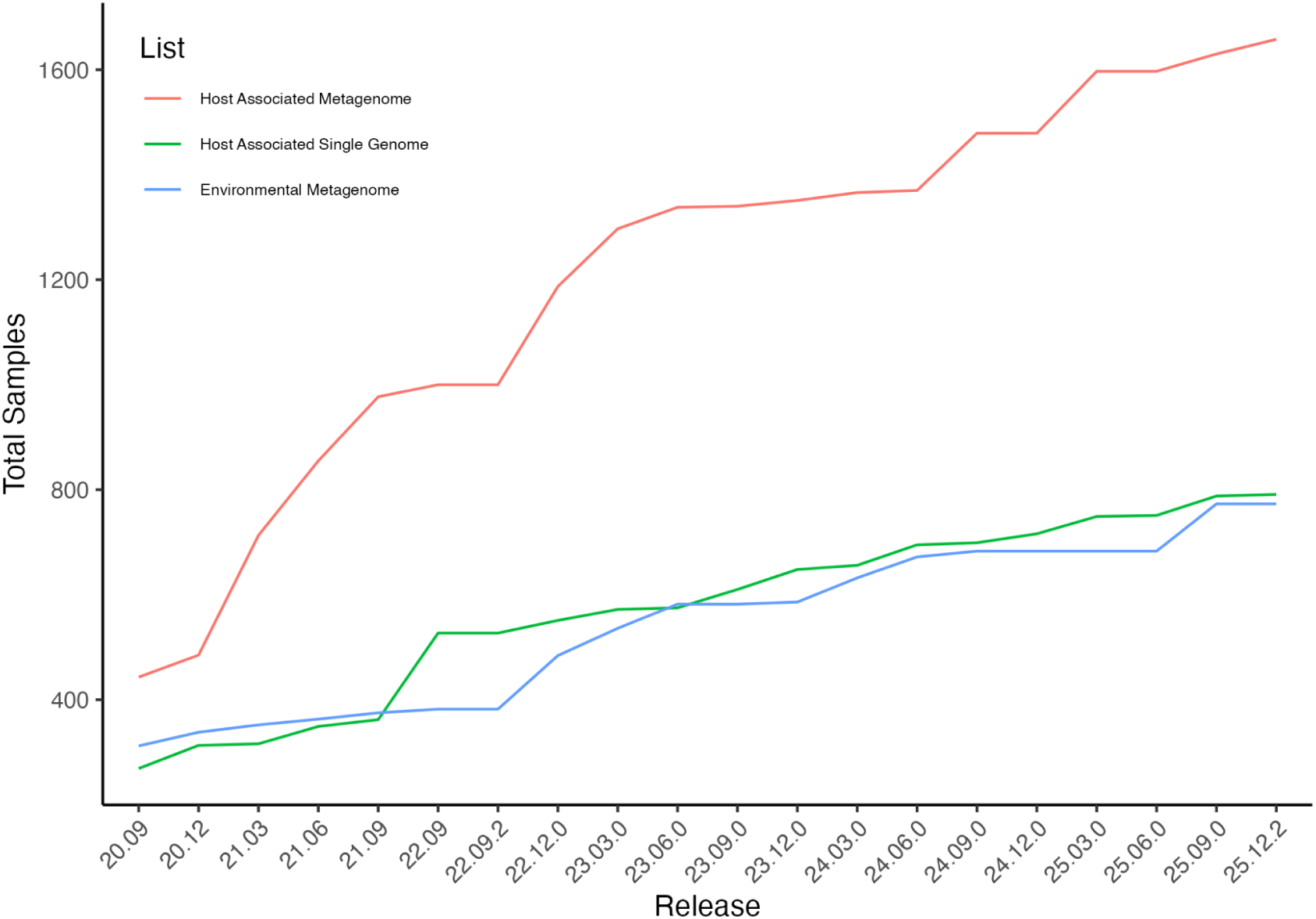
Cumulative number of samples included in each release of AncientMetagenomeDir. Since its initial release (20.09), the number of samples included has increased for ancient metagenomes associated with a host from 443 to 1,658, for ancient microbial genomes from 269 to 791, and for ancient environmental samples from 312 to 773, including metadata for a total of **3,222 samples**.

One limitation of early AncientMetagenomeDir releases was the reporting of dating information for a given sample as rounded median (to 100 years BP) of the primary date, without differentiation between absolute and relative dating methods. This simplified approach was adopted as a practical solution to the absence of standardised reporting guidelines for dates, and because extracting consistent and reliable dating information from publications proved challenging during the initial stages of building the AncientMetagenomeDir tables. When encountering radiocarbon dates, these challenges included inconsistencies in reporting uncalibrated radiocarbon dates vs calibrated dates (where the latter may be reported without the original uncalibrated date), and use of ranges or medians, requiring rounding or the addition of an extra field in the AncientMetagenomeDir. However, the use of a rounded date limits utilisation of the metadata set by researchers as the original estimates already include a degree of uncertainty, therefore is insufficient for accurate molecular dating and historical contextualization. Further approximating or rounding these dates can accumulate and amplify this uncertainty, thereby reducing the reliability of downstream analyses^28^.

Here, we present the first integration of standardised dating information into AncientMetagenomeDir, representing a collaborative project between aDNA researchers, archaeologists, and radiocarbon specialists. It includes precise dating information for each sample listed in release 25.09 of AncientMetagenomeDir’s *ancientsinglegenome-hostassociated* list that corresponds to ancient microbial genome sequencing data. Additionally, the information we gathered for the AncientMetagenomeDir dates extension allows us to quantitatively assess the prevalence of radiocarbon date reporting compliance within ancient metagenomics against conventions set by the radiocarbon community. We show a highly variable degree of reporting of radiocarbon dates that rarely fulfills the standards proposed by the radiocarbon community. This highlights the need for the introduction of a common, standardised, computer-readable and validatable file format and domain-neutral databases accepted by both the radiocarbon and palaeo– and archaeo-science community for archiving essential metadata of a given radiocarbon date.

## Methods

### Repository structure

AncientMetagenomeDir (https://doi.org/10.5281/zenodo.3980833) is a community-curated dataset maintained on GitHub (https://github.com/SPAAM-community/AncientMetagenomeDir) containing metadata from published ancient metagenomic studies. For the dating information, we adopted the same structure as the existing samples and libraries tables, using overlapping key columns to ensure that entries link easily with the other metadata in the repository. We systematically collected dates manually from all publications included in AncientMetagenomeDir’s *ancientsinglegenome-hostassociated* table (corresponding to ancient microbial genomes), and submissions to the table followed an updated version of the established AncientMetagenomeDir GitHub workflow (Figure 2).

**Figure 2.**
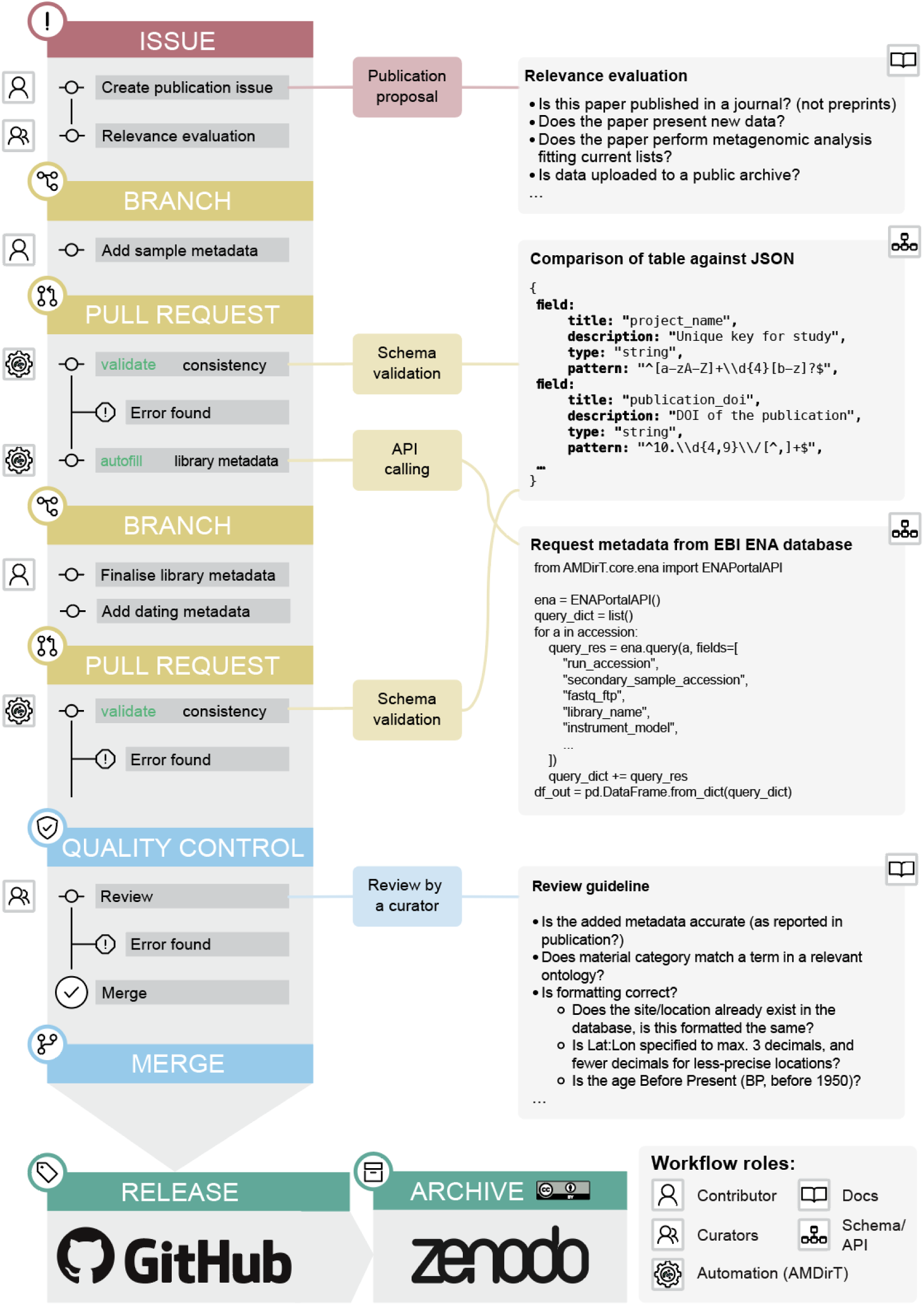
Updated AncientMetagenomeDir submission workflow with dating metadata information. Submissions are carried out on GitHub, and final releases are archived at Zenodo. Currently, “Add dating metadata” under the “Branch” section only applies to the entry of ancient microbial genome sequences *(ancientsinglegenome-hostassociated*) into AncientMetagenomeDir. Validation occurs with the same tooling as the existing sample and library metadata using the AMDirT toolkit.

Retaining the same structure and workflow benefits both experienced and new users, as experienced users are not required to learn a new structure or process, and new users only need to learn a single workflow to enter data into all three tables. Like the primary sample and library metadata lists in AncientMetagenomeDir, the dates extension table is formatted as a tab-separated value (TSV) file to allow interoperability between workflows and facilitate reusability for all researchers. The addition of sample dating information to AncientMetagenomeDir provides the benefit of the quality control assurances built into the existing AncientMetagenomeDir submission workflow using the associated AMDirT toolkit^26^, alongside providing easy access to public archive accession codes that guide researchers to associated sequence data.

### Data acquisition

The AncientMetagenomeDir dates extension has been built through two international community-sourced hackathon events attended by the authors, and through additional data entry by the main authors (K.H., D.S., A.A.V). For each publication in the AncientMetagenomeDir *ancientsinglegenome-hostassociated* table we opened an issue in the GitHub repository. A member of the SPAAM community then assigned themselves to the issue, acting as a contributor. The contributor created a Git branch from the main repository, manually retrieved the relevant metadata from the specified publication, and inserted it into the designated table following the specifications recorded in the repository’s documentation (https://www.spaam-community.org/AncientMetagenomeDir/#/docs/reference/). The introduced metadata was validated and subsequently reviewed by another member of the SPAAM community (see section “Technical Validation”). Additionally, all entries into the dates table are logged in the repository CHANGELOG, which effectively tracks the publications included in each release and can be referred to by a user, including any corrections. Documentation on how to submit a publication and sample dates data is available through tutorial documents accessible via the project website (https://www.spaam-community.org/AncientMetagenomeDir/#/docs/contributing/tutorials). In studies where the date information was unclear (e.g. if it was not clear whether the date was reported as calibrated or uncalibrated), we attempted to contact the original authors of the publication. These problematic studies (n=7, including 79 samples) remain excluded from the database until clarification from the authors is received.

## Data record

In the current release of AncientMetagenomeDir (v25.12.2: Historic Centre of Sighișoara https://github.com/SPAAM-community/AncientMetagenomeDir, archived at 10.5281/zenodo.18474769), dating information for ancient microbial samples

(*ancientsinglegenome-hostassociated*) was included due to immediate need by the community for this information in established analysis workflows, such as phylogenetic tip dating with BEAST, which is regularly performed in ancient microbial genome studies^29,30^.

The metadata in the *ancientsinglegenome-hostassociated_dates* table covers seven main categories: publication metadata, sample metadata, dating metadata, radiocarbon dating information, quality control information, calibration information, and reservoir offset information (Table 1). The fields included in the new dates schema were selected after discussion and consultation with ancient metagenomics researchers from the SPAAM community, archaeologists, and radiocarbon specialists. The basic column structure is based on the reporting conventions described in Millard (2014), with additional columns included to evaluate the quality of radiocarbon reporting and to allow inclusion of metadata for relative dating methods.

**Table 1:**
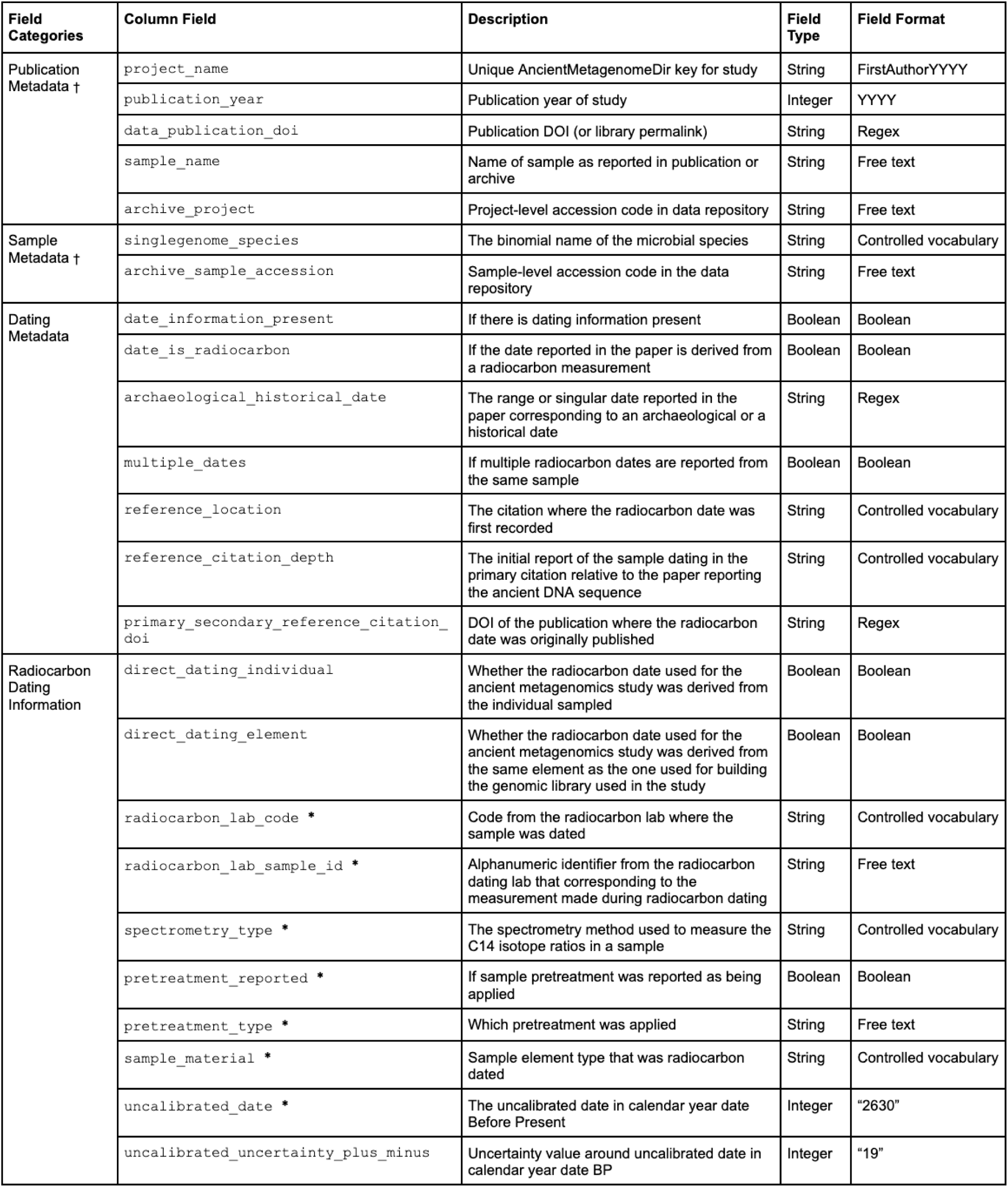
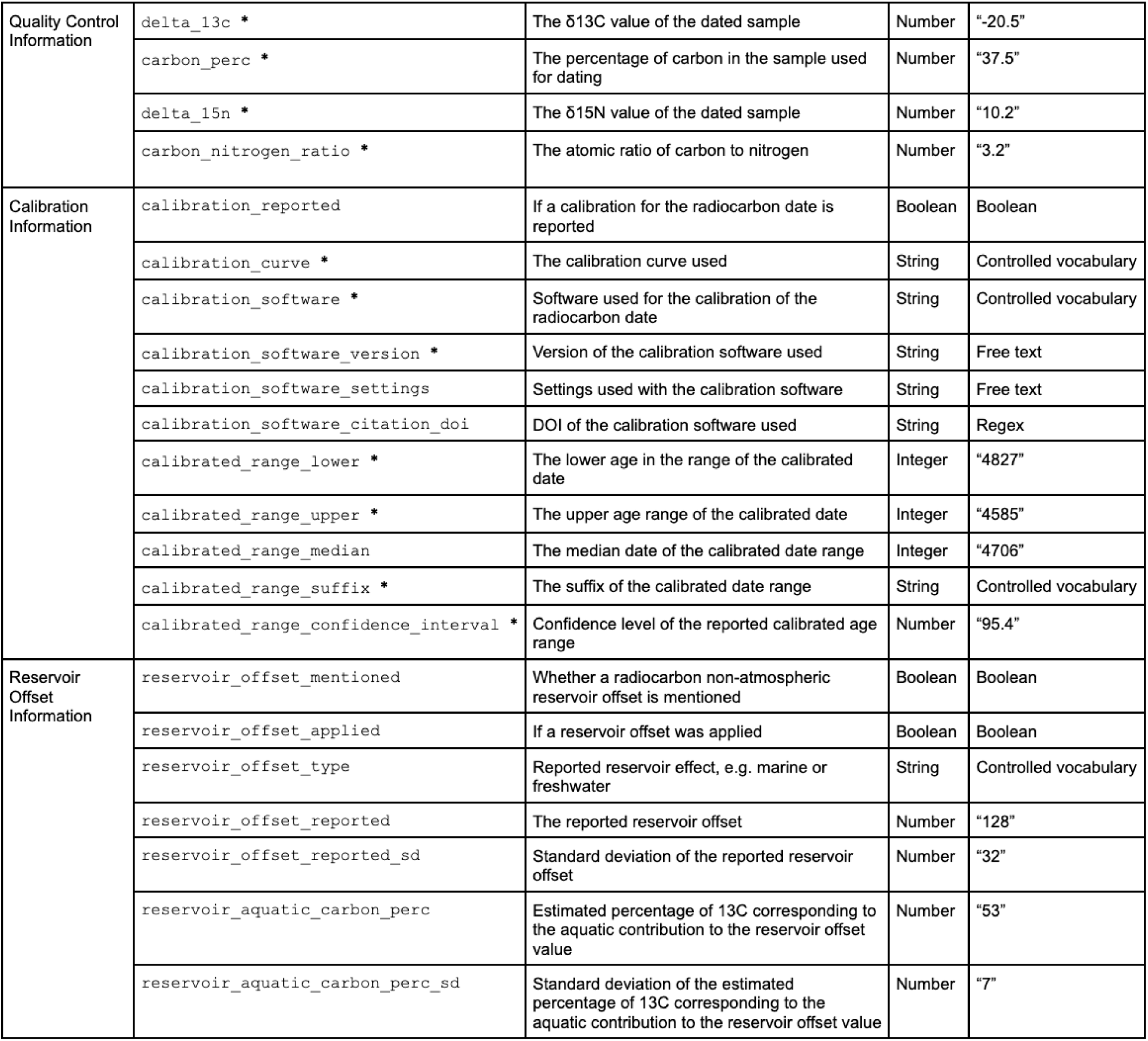
Complete Schema for dates table. Column fields marked with * are columns that are in accordance with the C14 reporting conventions described in Millard (2014) (supplementary note 1). Field categories marked with † are shared between the dates table and the sample and library tables to allow for cross-linking between the different tables in AncientMetagenomeDir.

The dates table includes dating information for all samples in the *ancientsinglegenome-hostassociated* list, including historically dated samples (e.g. museum specimens, samples from marked graves) and archaeologically dated samples (dated through stratigraphic or other archaeological methods) in addition to radiocarbon-dated samples. These are indicated by the `date_is_radiocarbon` and the `archaeological_historical_date` columns. The dates in the former two categories have been converted to ‘Before Present’ (BP) notation to allow easy comparison with radiocarbon-dated samples. No further metadata about historical or archaeological dates is recorded here due to the variability of methods and the lack of reporting standards. For example, samples dated using stratigraphic or typological methods are generally indicated by a range, whereas a historically dated sample is more likely to be given as a single date, but occasionally is also provided as a range.

Although reporting of the uncalibrated date is the recommended standard^4^ to allow for date recalibration (see introduction), we also included calibrated radiocarbon date information in our dataset to account for publications that only report the calibrated values and to aid meta-analysis. Metadata on radiocarbon analysis pretreatment and quality control assurances were included as they are essential for evaluating the robustness of a date, and are required reporting standards^4,31^. A lack of reporting of pretreatment protocols for bone collagen samples has recently been flagged as an issue across quaternary environmental fossil vertebrate research, given that pretreatment directly affects the dating accuracy of samples from this period^6^.

We also decided to differentiate between the columns `direct_dating_individual` and `direct_dating_element`. In some instances, the archaeological material used for aDNA extraction and the material used for radiocarbon dating may not be the same; this is important to be aware of, as the archaeological sample may have been disarticulated when excavated (perhaps leading to misidentification of two individuals as one) and human bone collagen offset will vary between different skeletal elements due to different bone remodelling rates^32^. Where proxy dates are used (indicated by `direct_dating_individual` marked `FALSE`), these issues are more pertinent. Finally, information about potential reservoir effects was included, as this can have a significant effect on the calculation of a calendar date. Samples that have undergone a reservoir correction in publications using outdated curves (as of the current date, prior to 2020) must be recalibrated, and thus this information is highly relevant during re-analysis with phylogenetic and other date-based analyses.

In total, the current AncientMetagenomeDir release (v25.12.2: Historic Centre of Sighișoara https://github.com/SPAAM-community/AncientMetagenomeDir, archived at 10.5281/zenodo.18474769) of the *ancientsinglegenome-hostassociated* dates table contains 768 entries, of which 738 are unique dates (with duplicates coming from the presence of multiple pathogens in a single host, as indicated by the `singlegenome_species` column). Of these 738 dates, 333 are historically or archaeologically dated, and 405 have radiocarbon dating information associated with them (Figure 3). Sample age is distributed from 9300 BP to the present, with radiocarbon dating being more frequent in samples until 2000 BP, where we see a steep increase in archaeological_historical dates (Figure 3), as expected for historical samples. The dataset includes dating information from almost all continents with a sample bias towards Eurasia (Figure 4, Fellows Yates et al., 2021). Whether this is due to biases, such as access to funding and collections, or preservation, is out of the scope of the present study.

**Figure 3.**
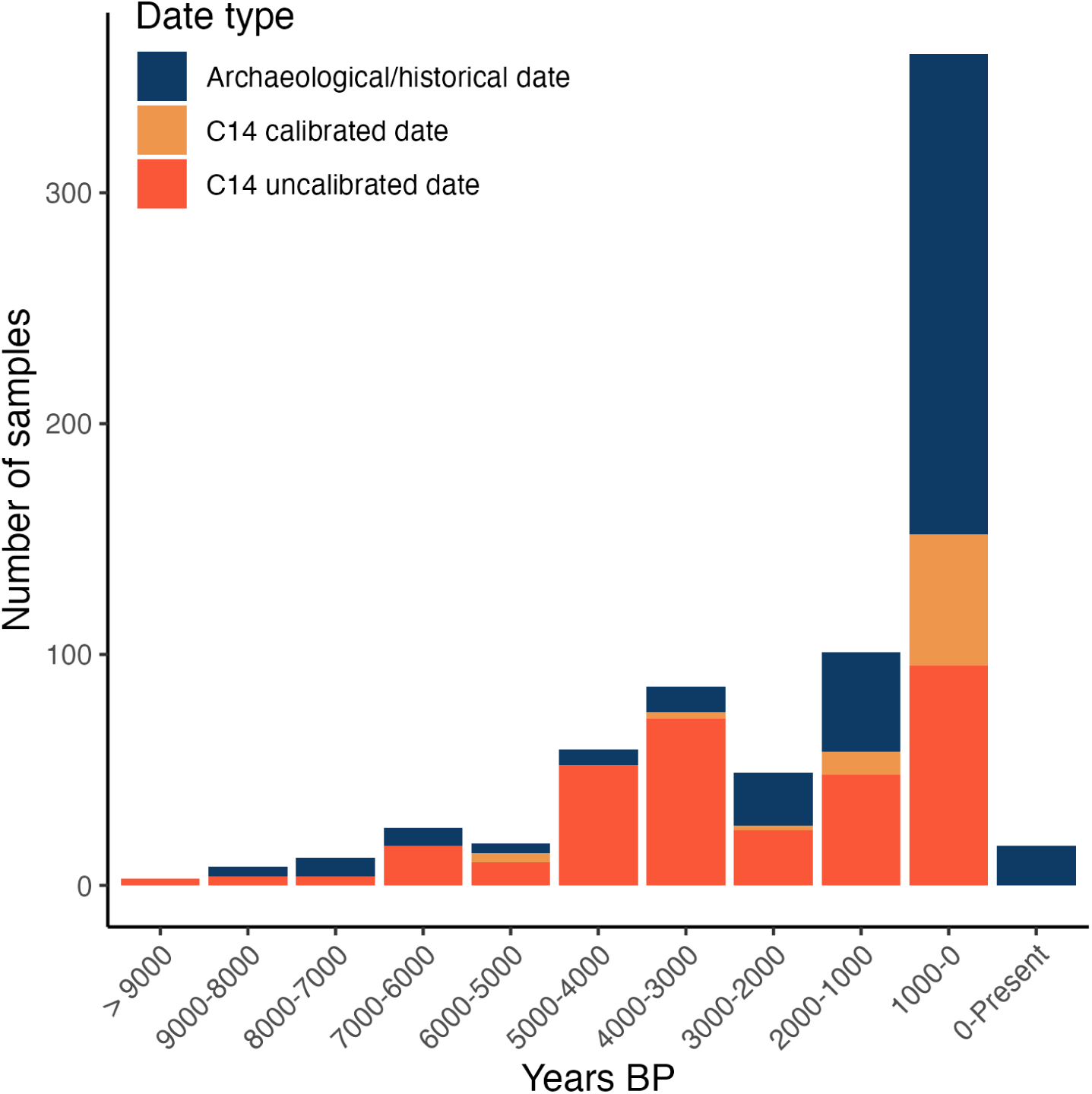
Date distribution of microbial samples grouped into 1,000-year intervals. Colours represent the type of date: archaeological/historical date (blue, n=333), C14 calibrated date (where sample has no associated uncalibrated date provided,light orange, n=76) and C14 uncalibrated date (deep orange, n=329).

**Figure 4:**
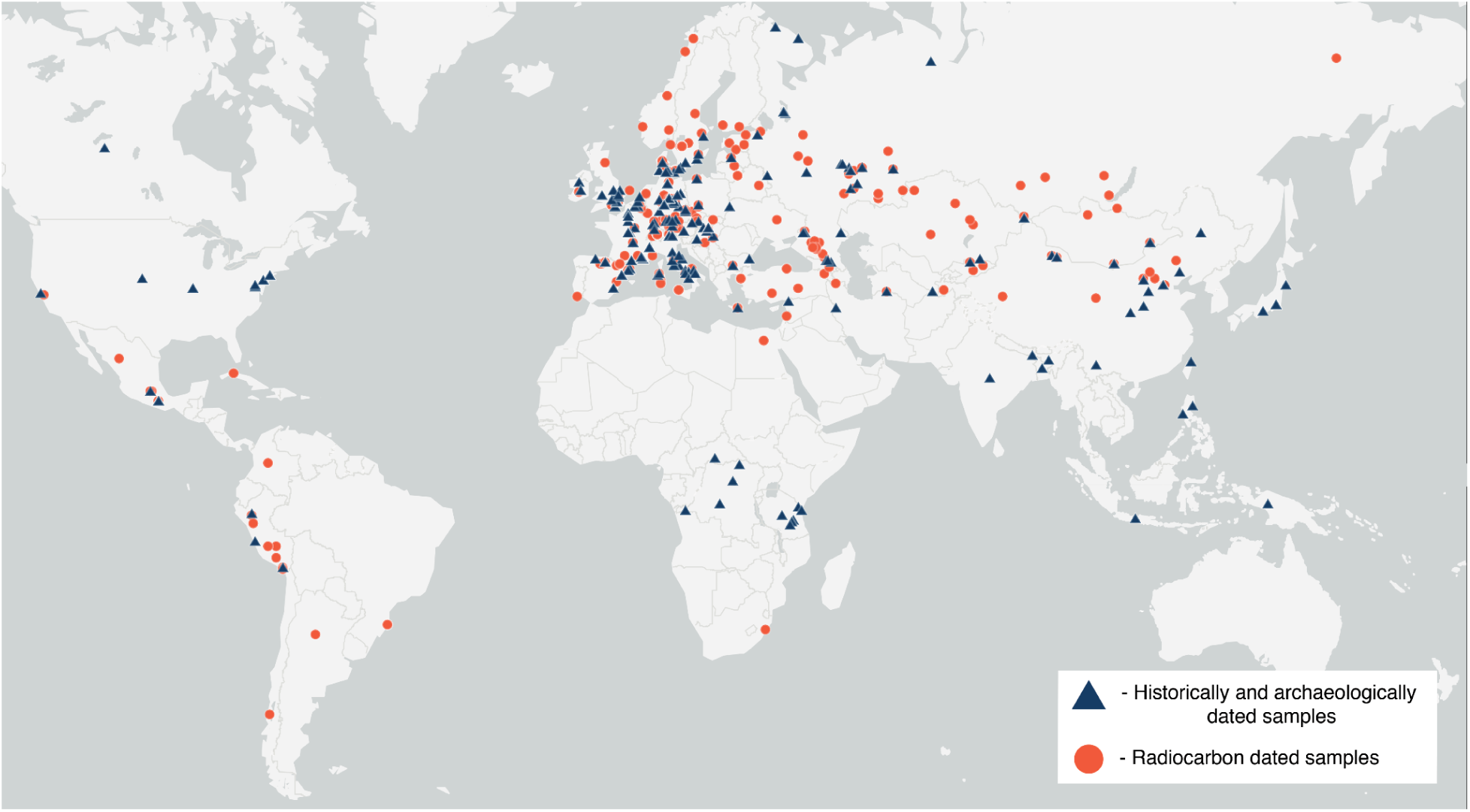
Geographical distribution of ancient microbial samples in the AncientMetagenomeDir dates extension. Generated using ArcGIS Online. Sources: Esri, TomTom, Garmin, FAO, NOAA, USGS, ©OpenStreetMap contributors, and the GIS User Community.

## Technical validation

During the addition of a date to the repository, a contributor initially assigns themselves to an issue within the AncientMetagenomeDir repository, retrieves the data, and updates the TSV file on a GitHub branch. Following the input of all metadata, the contributor then makes a Pull Request into the master branch in GitHub. Every Pull Request undergoes an automated continuous-integration validation check via the open-source tool AMDirT^26^; https://github.com/SPAAM-community/amdirt License: GNU GPLv3). These checks include assessing conformity against a JSON specification schema version of our written specifications, including format validation of specified fields (e.g. DOIs, project IDs, date formats), and the cross-checking of entries against controlled vocabularies defined in auxiliary JSON-based “enum” files^25^, with terms often derived from established term-ontologies. We also incorporated additional controlled vocabulary specific to the dating information. For example, the JSON schema corresponding to the `radiocarbon_lab_code` column follows the standardised abbreviations as maintained by the radiocarbon laboratory community through the *Radiocarbon* journal (https://radiocarbon.org/laboratories).

Following clearance of the automated checks, the contributor must request a peer review from another member of the SPAAM community. The reviewer checks for both the accuracy of the added metadata against the original publication and the consistency against the table’s README specification file. When these checks are complete, the metadata for the publication are merged with the master branch, and the corresponding Issue is closed.

AncientMetagenomeDir is maintained on GitHub and has regular quarterly releases, each with a release-specific DOI assigned via the Zenodo long-term data repository. From release v25.06, the database included data for the ancient microbial samples dates extension in a TSV format entitled *ancientsinglegenome_hostassociated_dates*.

## Usage Notes

Use of the AncientMetagenomeDir dates table extension typically requires the user to download the dataset from either GitHub or Zenodo. We have extended the command line interface of the accompanying tool AMDirT^26^ to facilitate download of the complete dates table (amdirt download –t ancientsinglegenome-hostassociated –y dates)and allow users to filter this table based on a prefiltered samples table (amdirt convert –-dates samples_table.tsv ancientsinglegenome-hostassociated), enabling the manipulation of the dates table without the need for additional software, in addition to producing BibTeX files with all the required literature for referencing. In the future, we plan to incorporate this functionality into the AMDirT ‘viewer’ graphical user interface. If the user downloads the dataset via GitHub or Zenodo, they must load the relevant TSV file into software such as Microsoft Excel, LibreOffice Calc, or R. The table can then be sorted, filtered, and queried according to user preference.

All dates have been added on a best-effort basis. Therefore, if a dated sample is used in an analysis, the user should independently check the date and associated metadata to evaluate the robustness of the sample date, such as whether a reservoir effect may be present which has not been corrected for, or whether an archaeological date is derived from a disputed context. Note that all data retrieved using AncientMetagenomeDir that is used in subsequent studies should be cited using the original publication citation, in addition to citing AncientMetagenomeDir.

## Discussion

By providing comprehensive and standardised dating information for ancient microbial genomes, the AncientMetagenomeDir dates extension further improves the ability of researchers across different palaeo– and archaeo-scientific disciplines to easily find and reuse publicly available comparative metagenomic data. Alongside the existing AncientMetagenomeDir geographic location metadata and publicly accessible sequence accession numbers, the precise dating information that can now be retrieved from this extension will allow researchers to more efficiently and robustly perform molecular dating and other evolutionary analyses in studies of ancient pathogen emergence and diversification.

To ensure accuracy the user is encouraged to recalibrate the given raw radiocarbon date(s) using the most recent calibration curve, ensuring consistency in precision between radiocarbon dates produced at different time points. To facilitate the re-use of the dates, we aim to implement automatic recalibration using an external recalibration software within AMDirT. With the framework now established for ancient microbial genomes, we are expanding dating information tables to the other represented sample types in future releases of AncientMetagenomeDir, as the generation of metagenomic assembled genomes (MAGs) is now becoming routine in both ancient microbiome and sedaDNA microbial studies^33–35^ and we expect these will also routinely be included in phylogenomic analyses. The *ancientmetagenome-environmental* sample list, however, will pose a different challenge, since dating of environmental samples, such as sediment, is achieved by a wider range of different absolute dating methods beyond radiocarbon dating. This makes standardisation more challenging, by requiring different standards for different dating methods, and an extended and more complex validation schema.

The process of aggregating and standardising dating information for samples in AncientMetagenomeDir has also provided us an opportunity to undertake a preliminary meta-analysis regarding C14 reporting convention compliance in the field of ancient metagenomics, specifically for studies focusing on the recovery of ancient microbial genomic sequences. To conduct this analysis, we checked each column based on the conventions described in ^4^ to calculate the proportion of missing dating information for current metagenomic datasets in the database.

There are currently 738 total samples in the AncientMetagenomeDir dates extension, of which 405 have C14 information (note: some samples have multiple measurements). Of these 405, none of the dates had 100% compliance with C14 reporting conventions as listed ^4^. The percentage completeness of each of these conventions for the samples in the AncientMetagenomeDir dates extension are presented in Table 2.

**Table 2:** Completeness of column data corresponding to C14 dating reporting conventions described in Millard (2014). All samples are of a proteinaceous type, with the exception of 2 wood samples.

| Field name | Sample size | Sample rows including data | Sample rows data not reported | metadat a present | not reported |
| --- | --- | --- | --- | --- | --- |
| radiocarbon_lab_code | 405 | 360 | 45 | 88.89% | 11.11% |
| radiocarbon_lab_sample_id | 405 | 315 | 90 | 78.78% | 22.22% |
| sample_material | 405 | 292 | 113 | 72.10% | 27.90% |
| pretreatment_type | 405 | 70 | 335 | 17.28% | 82.72% |
| uncalibrated_date | 405 | 329 | 76 | 81.23% | 18.77% |
| delta_13c | 405 | 99 | 306 | 24.44% | 75.56% |
| delta_15n | 405 | 61 | 344 | 15.06% | 84.94% |
| carbon_nitrogen_ratio | 405 | 68 | 337 | 16.79% | 83.21% |

One of the useful data reporting conventions agreed upon amongst the radiocarbon community is a community agreed set of standardised lab codes maintained by the journal *Radiocarbon*, which theoretically allows traceability of a date back to the original data-producing laboratory^4^. These alphanumeric codes consist of a sequence of letters, indicating the lab itself (lab ID), followed by a lab-specific date identifier (date ID), indicating the specific measurement of a sample^31^. For some labs that host online databases (e.g. the Oxford Radiocarbon Accelerator Unit online database, which is searchable by OxA date IDs), the complete lab code can provide access to additional sample metadata held by the radiocarbon lab. In the *ancientsinglegenome-hostassociated* dates list, there are 360 samples associated with lab codes, meaning for 45 samples (11.11%) included in our database, the radiocarbon lab codes are absent and therefore untraceable (Table 2). For 90 samples (22.22%), although the lab ID may be reported, the sample-specific dateID is absent, meaning radiocarbon date metadata for that particular sample cannot be traced. We can see a slight shift in these percentages if we exclude dates from secondary literature (hereby referring to cited publications containing information about the samples included in the AncientMetagenomeDir table, but not in the publication reporting the ancient metagenomic data itself); in primary metagenomics publications, date IDs are more commonly absent (Supplementary Table 1). The omission of lab code information is a significant issue because, even if the laboratory is known, retrievability is challenging given the large volume of data produced by many radiocarbon facilities.

Our dataset contains 405 radiocarbon dates, however uncalibrated dates are missing for 76 samples (Table 2). This is problematic, because unreported raw radiocarbon dates prevent re-calibration with updated calibration curves. Scientists researching ancient microbial evolution require accurately calibrated sample ages to serve as reliable tip dates for phylogenomic analysis^29^. Re-calibration of radiocarbon dates can sufficiently alter the outcome of evolutionary analyses as to shift potential interpretation of ancient pathogen dynamics, affecting the precision of analysis using programmes such as BEAST^36^.

In an analysis of a dataset of over 100,000 radiocarbon dates from Holarctic large-bodied mammal collagen, Herrando-Pérez and Stafford (2025) shed light on data curatorial shortfalls, in which they found that unpublished pretreatment information (50% in their dataset) was often challenging to trace when laboratories were directly approached, emphasising the importance of effective data reporting at the outset. Our dataset yielded significantly lower rates of reporting of pretreatment (82.72% absence) in metadata associated with ancient microbial data, preventing future researchers from evaluating the robustness of previously generated radiocarbon dates.

We also found that measurements used for ^14^C quality assurance and non-atmospheric offset detection had data absence rates ranging from 75-85% (e.g., C:N ratio (`carbon_nitrogen_ratio`), δ13C (`delta_13c`), δ15N (`delta_15n`)). As a third of the dates in our table come from secondary or tertiary publications, we evaluated whether the standards in reporting are similarly lacking in publications conducting primary dating analysis for individuals that yielded ancient microbial genome sequences (Supplementary Table 1). Our data suggests a higher prevalence of reporting of quality control values in primary ancient microbial genome publications (representing ‘new’ C14 dates) than secondary or tertiary citations. However, the widespread absence of reporting of quality control values across both primary metagenomic and secondary and tertiary palaeogenomic and archaeological literature prevents researchers from independently assessing data quality. While it can be assumed that a researcher would not publish a radiocarbon date falling below quality metrics, it may be that future researchers have different quality thresholds. As two thirds of the C14 dates in our dataset represent the first publication of sample C14 dating information insufficient metadata provision prevents re-use of this data by all future researchers.

New radiocarbon dating information is increasingly produced in the context of aDNA studies, therefore, adequate radiocarbon metadata reporting is increasingly becoming an additional responsibility of palaeogenomicists themselves. As such, good reporting standards are imperative to maximise the value to the wider scientific community. Given this responsibility, in a field celebrated for its excellent data reporting practices^23^, we hypothesise reasons for the insufficient reporting of radiocarbon dating information. One reason may be a lack of training and understanding of radiocarbon dating methodology. This gap in understanding between radiocarbon specialists and other archaeological scientists is a widespread issue, and radiocarbon dating has been described as being seen as a “black box”^7^. Therefore, researchers reporting C14 dating information may not always have the knowledge or understand the necessity in publishing comprehensive metadata in standard formats. Given our evaluation of radiocarbon data reporting in ancient metagenomic studies, we also want to raise awareness among peer reviewers to pay attention to this issue, and promote minimum reporting standards (e.g., lab IDs) such as those proposed in Millard (2014), where data reporting is insufficient in initial manuscripts.

Finally, for a predominantly data– and bioinformatic centered field that is accustomed to community-accepted standards, such as FASTQ^21^ or BAM files^22^, the lack of a well-defined file format for the reporting of radiocarbon dating information is likely restricting the effective dissemination of this data. An accepted common, standardised, computer-readable and validatable file format originating from the radiocarbon community (in collaboration with palaeogenomics and other areas of archaeological sciences) would be an important step towards addressing the inconsistent reporting of radiocarbon dating information for ancient metagenomics studies, and likely in archaeology more widely. Such a standard would also facilitate the future creation of a generalised and centralised database for all C14 dates akin to the popular ENA/SRA within genomics (as also called for in ^6^, and increase the ability to link metadata between many different sources of archaeological information through stable identifiers.

We recognise that the insufficient publication of essential radiocarbon dating information is not limited to ancient metagenomics, nor is it specific to the field of aDNA. In fact, the publication of dates and associated metadata was described over a decade ago as the “largest and most pressing problem facing the field” of radiocarbon research. This problem persists today, and as datasets grow in size, addressing it becomes increasingly critical^6,8^. As ancient metagenomics researchers, we can continue to sustain the precedent of high data curatorial standards in palaeogenomics, and expand our commitment to open data sharing^23^ to include dating information. The two initiatives we have suggested: the introduction of a community-accepted common, standardised, computer-readable and validatable file format; and the development of a centralised database for all C14 dates, will benefit all archaeological sciences. The creation of the AncientMetagenomeDir dates extension is an initial step in this direction.

## Data availability

The most up to date version of the dataset can be found in the AncientMetagenomeDir GitHub repository: https://github.com/SPAAM-community/AncientMetagenomeDir. All versions of this dataset are archived in Zenodo under DOI: 10.5281/zenodo.3980833. The version presented in this article is v25.12.2: Historic Centre of Sighișoara, which can also be accessed in Zenodo DOI: 10.5281/zenodo.18474769.

## Code availability

An R notebook used for generating images with package versions can be found in the AncientMetagenomeDir repository at https://github.com/SPAAM-community/AncientMetagenomeDir/tree/master/assets/analysis/ (commit 501bb99).

Code for validation of the dataset can be found at https://github.com/SPAAM-community/amdirt and https://doi.org/10.5281/zenodo.4003826.

## Author Contributions

Conceptualization: J.A.F.Y., A.A.V., K.H, D.S. Metadata collection: All coauthors. Data curation: A.A.V., K.H., J.A.F.Y., D.S., B.P.B., B.B., Y.H., A.H., P.R. Data analysis of radiocarbon date reporting: K.H., A.A.V. Writing – original draft: K.H., A.A.V., J.A.F.Y. Writing – review & editing: All co-authors.

## Competing Interests

The authors declare no competing interests.

## Acknowledgements

We thank Silvia Tardaguila Giacomozzi and Anan Ibrahim for their input and help collecting some metadata; Amy Bogaard and Greger Larson for K.H. supervision, Rachel Wood for comments. We thank Alexander Herbig, Johannes Krause, and the SPAAM community for their support in this project.

## Funding

The authors acknowledge funding by the Max Planck Society (A.A.V., J.A.F.Y, I.J.), Merton College Graduate Archaeology Scholarship (K.H.), the Clarendon Fund (K.H.), SciLifeLab and Wallenberg Data Driven Life Science Program (KAW 2020.0239, C.J.); European Research Council (ERC) under the European Union’s Horizon 2020 Research and Innovation Programme (SEACHANGE, grant agreement no. 856488, G.Z.); Leverhulme Trust Fund Early Career Research Fellowship (ECF-2022-532, to L.B.V.); Swiss National Science Foundation Ambizione grant PZ00P1_223787 (M.K.); Social Sciences and Humanities Research Council [Doctoral – Vanier Canada Graduate Scholarship, 2024], Pierre Elliott Trudeau Foundation [Scholar, 2024], and Simon Fraser University (C.N.H.T.); Deutsche Forschungsgemeinschaft (DFG, German Research Foundation) under Germanýs Excellence Strategy (EXC 2051 – Project-ID 390713860, “Balance of the Microverse”, A.H., C.W.); Deutsche Forschungsgemeinschaft (DFG, German Research Foundation) – 460129525 (NFDI4Microbiota FlexFund: “EnterArchaeo”, J.A.F.Y., C.W.); Werner Siemens Stiftung (“Paleobiotechnology”, I.V., C.W., J.A.F.Y.); Carolyn Weinberg and the Radcliffe Institute for Advanced Study (C.W.).

## Notes

https://zenodo.org/records/18474769

### Competing Interest Statement

The authors have declared no competing interest.

## References

1. Hajdas, I. et al. Radiocarbon dating. Nat. Rev. Methods Primers 1, (2021).

2. Libby, W. F., Anderson, E. C. & Arnold, J. R. Age Determination by Radiocarbon Content: World-Wide Assay of Natural Radiocarbon. Science 109, 227–228 (1949).

3. Arnold, J. R. & Libby, W. F. Age determinations by radiocarbon content; checks with samples of known age. Science 110, 678–680 (1949).

4. Millard, A. R. Conventions for reporting radiocarbon determinations. Radiocarbon 56, 555–559 (2014).

5. Bayliss, A. Quality in Bayesian chronological models in archaeology. World Archaeol. 47, 677–700 (2015).

6. Herrando-Pérez, S. & Stafford, T. W., Jr. Making vertebrate fossil radiocarbon dates more useful for global scientific research. J. Quat. Sci. (2025) doi:10.1002/jqs.70012.

7. Wood, R. From revolution to convention: the past, present and future of radiocarbon dating. J. Archaeol. Sci. 56, 61–72 (2015).

8. Becerra-Valdivia, L. Climate influence on the early human occupation of South America during the late Pleistocene. Nat. Commun. 16, 2780 (2025).

9. Brace, S. et al. Ancient genomes indicate population replacement in Early Neolithic Britain. *Nat*. Ecol. Evol. 3, 765–771 (2019).

10. Moreno-Mayar, J. V. et al. Ancient Rapanui genomes reveal resilience and pre-European contact with the Americas. Nature 633, 389–397 (2024).

11. Yaka, R. et al. Variable kinship patterns in Neolithic Anatolia revealed by ancient genomes. Curr. Biol. 31, 2455–2468.e18 (2021).

12. Rivollat, M. et al. Extensive pedigrees reveal the social organization of a Neolithic community. Nature 620, 600–606 (2023).

13. Kocher, A. et al. Ten millennia of hepatitis B virus evolution. Science 374, 182–188 (2021).

14. Seersholm, F. V. et al. Repeated plague infections across six generations of Neolithic Farmers. Nature 632, 114–121 (2024).

15. L’Hôte, L. et al. An 8000 years old genome reveals the Neolithic origin of the zoonosis Brucella melitensis. Nat. Commun. 15, 6132 (2024).

16. Guinet, B. et al. Ancient host-associated microbes obtained from mammoth remains. Cell (2025) doi:10.1016/j.cell.2025.08.003.

17. Sikora, M. et al. The spatiotemporal distribution of human pathogens in ancient Eurasia. Nature 643, 1011–1019 (2025).

18. Ekram, M. A.-E. et al. A 1 Ma sedimentary ancient DNA (sedaDNA) record of catchment vegetation changes and the developmental history of tropical Lake Towuti (Sulawesi, Indonesia). Geobiology 22, e12599 (2024).

19. Seersholm, F. V. et al. Rapid range shifts and megafaunal extinctions associated with late Pleistocene climate change. Nat. Commun. 11, 2770 (2020).

20. Rijal, D. P. et al. Sedimentary ancient DNA shows terrestrial plant richness continuously increased over the Holocene in northern Fennoscandia. Sci. Adv. 7, eabf9557 (2021).

21. Cock, P. J. A., Fields, C. J., Goto, N., Heuer, M. L. & Rice, P. M. The Sanger FASTQ file format for sequences with quality scores, and the Solexa/Illumina FASTQ variants. Nucleic Acids Res. 38, 1767–1771 (2010).

22. Li, H. et al. The Sequence Alignment/Map format and SAMtools. Bioinformatics 25, 2078–2079 (2009).

23. Anagnostou, P. et al. When data sharing gets close to 100%: what human paleogenetics can teach the open science movement. PLoS One 10, e0121409 (2015).

24. Lien-Talks, A. How FAIR is bioarchaeological data: With a particular emphasis on making archaeological science data reusable. J. Comput. Appl. Archaeol. 7, 246–261 (2024).

25. Fellows Yates, J. A., et al. Community-curated and standardised metadata of published ancient metagenomic samples with AncientMetagenomeDir. Sci. Data 8, 31 (2021).

26. Borry, M. et al. Facilitating accessible, rapid, and appropriate processing of ancient metagenomic data with AMDirT. F1000Res. 12, 926 (2023).

27. Wilkinson, M. D. et al. The FAIR Guiding Principles for scientific data management and stewardship. Sci. Data 3, 160018 (2016).

28. Dos Reis, M. & Yang, Z. The unbearable uncertainty of Bayesian divergence time estimation: Uncertainty in divergence time estimation. J. Syst. Evol. 51, 30–43 (2013).

29. Bos, K. I. et al. Paleomicrobiology: Diagnosis and evolution of ancient pathogens. Annu. Rev. Microbiol. 73, 639–666 (2019).

30. Baele, G. et al. BEAST X for Bayesian phylogenetic, phylogeographic and phylodynamic inference. Nat. Methods 22, 1653–1656 (2025).

31. Stuiver, M. & Polach, H. A. Discussion: reporting of 14C data. Radiocarbon 19, 355–363 (1977).

32. Quinn, R. L., Beasley, M. M., Gocha, T. P. & Mavroudas, S. R. Differential human bone remodeling rates and implications for the temporal resolution of geoprofiling isotopes. Forensic Sci. Int. 370, 112454 (2025).

33. Wibowo, M. C. et al. Reconstruction of ancient microbial genomes from the human gut. Nature 594, 234–239 (2021).

34. Klapper, M. et al. Natural products from reconstructed bacterial genomes of the Middle and Upper Paleolithic. Science 380, 619–624 (2023).

35. Zhong, M. et al. Climate-driven deoxygenation promoted potential mercury methylators in the past Black Sea water column. *Nat*. Water 1–8 (2025).

36. Suchard, M. A. et al. Bayesian phylogenetic and phylodynamic data integration using BEAST 1.10. Virus Evol. 4, vey016 (2018).

