## Supplementary Note 1 & Table 1 for "AncientMetagenomeDir dating metadataset highlights need for standardised radiocarbon reporting in ancient DNA"

[1] Institute of Archaeology, University of Oxford, Oxford, U.K.; [2] Independent Researcher; [3] Department of Archaeogenetics, Max Planck Institute for Evolutionary Anthropology, Leipzig, Germany; [4] Associated Research Group of Archaeogenetics, Leibniz Institute for Natural Product Research and Infection Biology – Hans Knöll Institute, Jena, Germany; [5] Estonian Biocentre, Institute of Genomics, University of Tartu, Tartu, Estonia; [6] Globe Institute, University of Copenhagen, Copenhagen, Denmark; [7] Mechanical Engineering, Delft University of Technology, Delft, Netherlands; [8] Archaeological Sciences, Leiden University, Leiden, Netherlands; [9] Institute of Biodiversity, Ecology, and Evolution, Faculty of Biological Sciences, Friedrich Schiller University, Jena, Germany; [10] Department of Anthropology and Archaeology, University of Bristol, Bristol, U.K.; [11] Department of Computational Biology, University of Lausanne, Lausanne, Switzerland; [12] Swiss Institute of Bioinformatics, Lausanne, Switzerland; [13] Ancient Genomics Laboratory, The Francis Crick Institute, London, U.K.; [14] School of Biosciences, Cardiff University, Cardiff, U.K.; [15] Natural History Museum, London, U.K.; [16] The Gray Faculty of Medical & Health Sciences, Tel Aviv University, Tel Aviv, Israel; [17] The Dan David Center for Human Evolution and Biohistory Research, Tel Aviv University, Tel Aviv, Israel; [18] Department of Genetics Evolution and Environment, UCL Genetics Institute, University College London, London, U.K.; [19] Pathogen evolution, Helmholtz-Institute for One Health, Greifswald, Germany; [20] Department of Bioinformatics and Genetics, Swedish Museum of Natural History, Stockholm, Sweden; [21] Institute of Clinical Molecular Biology, Kiel University, Kiel, Germany; [22] Department of Anthropology, Dartmouth College, Hanover, USA; [23] Thayer School of Engineering, Dartmouth College, Hanover, USA; [24] Centre for Palaeogenetics, Stockholm, Sweden; [25] Department of Zoology, Stockholm University, Stockholm, Sweden; [26] Department for Environmental Sciences, University of Basel, Basel, Switzerland; [27] Institute of Genomics, University of Tartu, Tartu, Estonia; [28] Transmission, Infection, Diversification & Evolution Group, Max Planck Institute of Geoanthropology, Jena, Germany; [29] Max Planck Institute for Infection Biology, Berlin, Germany; [30] Evolutionary Dynamics of Infectious Diseases Unit, Institut Pasteur, Université Paris Cité, Paris, France; [31] Department of Human Evolutionary Biology, Harvard University, Cambridge, USA; [32] University of Aveiro, Aveiro, Portugal; [33] Animal Ecology, Wageningen Environmental Research, Wageningen, Netherlands; [34] Department of History and Art History, Utrecht University, Utrecht, Netherlands; [35] Department of Archaeology and Classical Studies, Stockholm University, Stockholm, Sweden; [36] Instituto de Antropología de Córdoba CONICET-UNC, Córdoba, Argentina; [37] Departamento de Antropología, Facultad de Filosofía y Humanidades, Universidad Nacional de Córdoba, Córdoba, Argentina; [38] Microbial Paleogenomics Uni, Institut Pasteur, Université Paris Cité, CNRS UMR 2000, Paris, France; [39] Ancient DNA Lab, Institute of Molecular Biology and Biotechnology (IMBB), Foundation for Research and Technology – Hellas (FORTH), Irakleio, Crete, Greece; [40] Department of Archaeology, Simon Fraser University, Burnaby, Canada; [41] Department of Anthropology, Harvard University, Cambridge, USA; [42] Department of Palaeobiotechnology, Leibniz Institute for Natural Product Research and Infection Biology – Hans Knöll Institute, Jena, Germany

[*] Authors contributed equally
† Corresponding authors

### Supplementary Note 1

Millard's (2014) conventions for radiocarbon dating metadata and our corresponding columns are as follows:
"(1) the laboratory measurement should be reported as a conventional radiocarbon age in 14C yr BP or a fractionation-corrected fraction modern (F14C) value"
= `uncalibrated_date`
"(2) the laboratory code for the determination should be included"
= `radiocarbon_lab_code ; radiocarbon_lab_sample_id`
"(3) the sample material dated, the pretreatment method applied, and quality control measurements should be reported."
= `sample_material ; pretreatment_reported ; pretreatment_type ; delta_13c ; delta_15n ; carbon_nitrogen_ratio ; carbon_perc`
"(4) the calibration curve and any reservoir offset used"
= `calibration_curve ; reservoir_offset_mentioned ; reservoir_offset_type`
"(5) the software used for calibration, including version number, the options and/or models used, and wherever possible a citation of a published description of the software"
= `calibration_software ; calibration_software_version`
"(6) the calibrated date given as a range (or ranges) with an associated probability on a clearly identifiable calendar timescale."
= `calibrated_range_lower ; calibrated_range_upper ; calibrated_range_suffix ; calibrated_range_confidence_interval`

-

**Supplementary Table 1:** Percentage of radiocarbon dated samples that lack lab code and quality control reporting values in the AncientMetagenomeDir dates table. Primary: ancient metagenomic study in the AncientMetagenomeDir dataset; Secondary & Tertiary: publications about the samples in AncientMetagenomeDir that are not the ancient metagenomic study recorded in AncientMetagenomeDir.

| | **Primary** (271 total samples) | **Secondary & Tertiary** (134 total samples) | **Total missing** (405 total samples) |
|---|---|---|---|
| **Radiocarbon lab codes missing** | 11.07% (n= 30) | 11.19% (n=15) | 11.11 % (n = 45) |
| **Radiocarbon lab sample ID missing** | 27.67% (n=75) | 11.94% (n= 16) | 22.47% (n = 91) |
| **δ13C missing** | 71.21% (n = 193) | 84.33% (n = 113) | 75.55% (n = 306) |
| **δ15N missing** | 83.02% (n = 225) | 88.81% (n = 119) | 82.47% (n = 344) |
| **C:N missing** | 80.44% (n = 218) | 88.81 % (n = 119) | 83.21% (n = 337) |
| **Uncalibrated dates missing** | 24.35% (n = 66) | 7.46% (n = 10) | 18.91% (n = 76) |
